# Engineered full-length Fibronectin-based hydrogels sequester and present growth factors to promote regenerative responses in vitro and in vivo

**DOI:** 10.1101/687244

**Authors:** Sara Trujillo, Cristina Gonzalez-Garcia, Patricia Rico, Andrew Reid, James Windmill, Matthew J. Dalby, Manuel Salmeron-Sanchez

**Affiliations:** Centre for the Cellular Microenvironment, University of Glasgow, Glasgow, United Kingdom; Biomedical Research Networking Centre in Bioengineering biomaterials and Nanomedicine, Universitat Politècnica de València, Valencia, Spain; Centre for Ultrasonic Engineering, Department of Electronic and Electrical Engineering, University of Strathclyde, Glasgow, United Kingdom

## Abstract

Extracellular matrix (ECM)-derived matrices such as Matrigel are used to culture numerous cell types in vitro and can recapitulate certain ECM functions that support cell growth and differentiation. However, ECM-derived matrices suffer lot-to-lot variability, undefined composition and lack of controlled physical properties. There is a need to develop rationally designed synthetic matrices that can also recapitulate ECM roles. Synthetic matrices have certain limitations as they use synthetic peptides or fragments whereas the ECM consists of full proteins. Here, we report the development of degradable, PEG-based hydrogels of controlled stiffness that incorporate full-length fibronectin (FN) to enable solid-phase presentation of growth factors in a physiological manner. We demonstrate, in vitro and in vivo, the effect of incorporating vascular endothelial growth factor (VEGF) and bone morphogenetic protein 2 (BMP2), in these hydrogels to enhance angiogenesis and bone regeneration, respectively. We show that the solid-state presentation of growth factors enables very low growth factor doses to achieve regenerative effects.

## Introduction

Growth factors (GFs) are signalling molecules that play essential roles in tissue development and organogenesis. Some drive cell differentiation, while others stimulate cellular processes, such as migration or proliferation^1^. Given these roles, GFs have potential clinical utility, particularly in regenerative medicine^2,3^. Despite this, the potential of GFs has yet to be fully realised in translational medicine, partly due to their short half-life and rapid clearance in vivo^4,5^. Consequently, material-based strategies have been developed to control their sequestration and release, in order to reduce their required dosage and to control their local action^6–8^.

Bioinspired materials are a particularly interesting avenue of regenerative research. Such materials aim to exploit natural extracellular matrix (ECM) molecules because the ECM, and its component molecules, sequester GFs and facilitate their interactions with other ECM and cell-secreted molecules, to support regenerative cell processes. Thus, the ECM and its components can act as a GF reservoir and can coordinate the availability of GFs through ECM-GF interactions^9^. Glycosaminoglycans and proteoglycans are major components of the ECM that help to maintain its structure and can bind several GFs. For instance, the glycosaminoglycan heparin, has been incorporated into hydrogels to sequester GFs for tissue-engineering applications^10^. Other proteins, such as fibrinogen, tenascin C or fibronectin (FN) are also known to bind GFs^11–13^. In particular, FN is an ECM protein that binds a wide range of GFs, including therapeutically interesting GFs, such as bone morphogenetic protein 2 (BMP2, which drives bone formation^14^) and vascular endothelial GF (VEGF, which stimulates angiogenesis)^13^. FN presents its GF-binding site (FNIII_12-14_) next to a cell adhesion-binding site (FNIII_9-10_). The simultaneous binding of both by the cell triggers synergistic integrin/GF receptor signalling, which enhances the effect of GFs on stem cell differentiation and tissue repair and regeneration^11,15–18^.

Materials-based strategies have begun to emerge that exploit this synergistic interaction. Such approaches include 2D-surface strategies that target the efficient presentation of GFs by altering FN’s conformation. For example, poly(ethyl acrylate) (PEA) is a polymer that causes FN to spontaneously unfold into nano-networks. It has been used to expose the FNIII_9-10_ and FNIII_12-14_ binding sites to cells to enable the delivery of ultra-low doses of BMP2^19–21^. This PEA-based system has been used to present BMP2 in a murine non-healing (critical-size) bone defect model^22^ and in a veterinary case of a dog with a humeral fracture^21^. It has also been used to present VEGF in an in vivo wound healing murine model^20^. In 3D, fibrin hydrogels have been functionalised with FN fragments that contain either the cell adhesion binding site (FNIII_9-10_), the GF binding site (FNIII_12-14_) or both (FNIII_9-10_/FNIII_12-14_)^15^. These hydrogels, when loaded with both VEGF and platelet-derived GF (PDGF-BB), have been shown to promote wound healing in diabetic mice, and when loaded with BMP2, to promote bone regeneration in a rat non-healing bone defect model, compared to controls with either FNIII_9-10_ alone or FNIII_12-14_ alone ^15^.

However, the ECM is a 3D environment that contains full-length FN, rather than FN fragments or peptides. For instance, FN presents two binding sites (FNI_1-5_ and FNIII_12-14_) for proteoglycans, such as syndecan-2 and −4, which can trigger focal adhesion assembly via protein kinase C alpha (PKCα) and focal adhesion kinase (FAK) ^23,24^. Also, the variable region of FN contains the Leu-Asp-Val (LDV) and the Arg-Glu-Asp-Val (REDV) sequences that associate with α4β1 and α4β7 integrins (non Arg-Gly-Asp (RGD)-binding integrins)^25,26^. FN also contains FN-binding sites that are essential for FN fibrillogenesis (FNI_1-5_, FNIII_1-2_, FNIII_7_, FNIII_10_ and FNIII_12-15)_, which has been shown to contribute to vascular morphogenesis^27,28^. FN also has binding affinity for collagen and fibrin, among other molecules, demonstrating the versatile role of FN within the ECM^29–31^.

The ability to mimic the role of the ECM in GF presentation is key to developing effective materials-based approaches for regenerative medicine. The physiological, solid-state presentation of GFs is also required as their soluble delivery has significant limitations. For example, with soluble delivery, supraphysiological doses are required due to the short half-life of unbound GFs and to their clearance from the site. BMP2 represents an instructive example of the issues that can arise with traditional GF administration. The use of recombinant human (rh)BMP2 was approved by the Food and Drug Administration (FDA) in 2002; since then, BMP2 has been used as a clinical alternative for bone autografts, revolutionising the bone graft substitute market due to its highly osteo-inductive properties^32^. For instance, in the US between 2002 and 2007, rhBMP2 was used in nearly 25% of all spine fusion procedures and in 50% of all primary anterior lumbar interbody fusion procedures^33^. Currently, the concentration of BMP2 approved by the FDA is 1.5 mg/mL for human use. This high dose has led to the increased incidence of adverse effects, including inflammation, ectopic bone and tumour formation, wound and urogenital complications, and osteoclast activation^34^. Adverse effects have arisen from the clinical use of other GFs as well, such as with becaplermin (Regranex), which contains PDGF-BB, supraphysiological doses of which have been used, raising safety concerns^35^.

Bone regeneration is the second most-transplanted tissue after blood^36^. With bone autografts in short supply and with allografts being poorly bioactive^37,38^, angiogenesis is also of paramount importance to the health and survival of new or regenerating tissue^39^. In fact, both angiogenesis and osteogenesis are vital processes in acute fracture healing and bone repair. There is a significant crosstalk between these two signalling pathways^40,41^ and stem cells play key roles in both processes^42,43^. For example, mesenchymal stem cells (MSCs) are a major VEGF producer during vascular bone regeneration^44,45^. VEGF plays an integral role in the regulation of angiogenic pathways^46^, determining the pace of vascularisation^46,47^. The sequestration, delivery and presentation of VEGF has been engineered in 3D hydrogel constructs to promote endothelial cell activation and the formation of multicellular tube-like structures via either vasculogenic or angiogenic processes^48–51^. This bioactivity could be improved by producing hydrogels with which VEGF can interact with the FNIII_12-14_ region^52^.

To provide an alternative to natural ECM-derived matrices and to overcome some of the reported limitations of synthetic ones, we report here the development of synthetic, 3D hydrogels. These hydrogels contain full length, human FN, enabling ultra-low dose, solid-sate GF presentation. We also demonstrate that the physical properties of this hydrogel can be tuned to recapitulate the properties of native ECM. Using BMP2 and VEGF, we demonstrate the ability of this hydrogel system to recruit and retain GFs in vitro, thereby providing a novel 3D, functional environments with the potential to replace Matrigel, and to promote bone regeneration and vascularisation in vivo.

## Results

### Fabrication and characterisation of full-length FN hydrogels

FN was covalently incorporated into a poly(ethylene) glycol (PEG) network via Michael-type addition reaction, where a 4-arm-PEG-maleimide spontaneously reacts with a thiolated crosslinker at physiological pH (Figure 1a, b). Maleimide groups have higher affinity towards thiol groups, which confer shorter gelation times at physiological pH relative to hydrogels fabricated using acrylate groups^53^. Michael-type addition has been previously used to form biologically active and cytocompatible hydrogels^54,55^.

**Figure 1.**
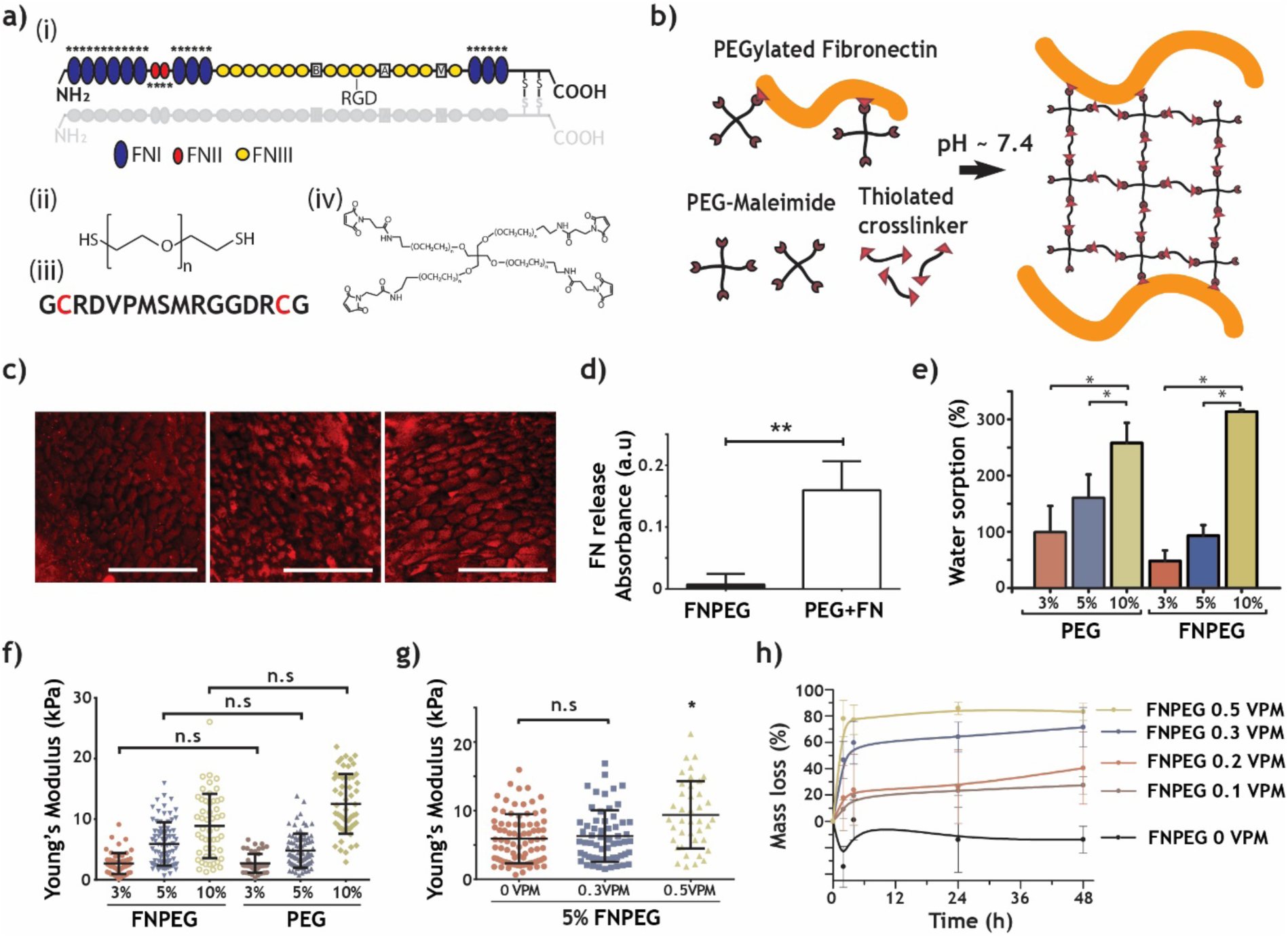
FN hydrogel formation and characterisation. **(a)** (i) Schematic of the modular composition of FN (upper structure, domains I, II and III are depicted in blue, red and yellow, respectively, and cysteine residues are marked as *), the crosslinker (SH-PEG-SH (ii) and degradable “VPM” peptide flanked by two cysteine residues (depicted in red) (iii)) and of 4-arm-PEG-maleimide (iv). **(b)** A schematic of the hydrogel formation protocol, where FN is depicted in orange. **(c)** Immunofluorescence of FN (in red) in cryosections of 3, 5 and 10 wt. % hydrogels (from left to right, scale bar: 100 µm). **(d)** FN release (normalised to PEG-only hydrogels) (mean ±SD, n = 3) from hydrogels into which PEGylated FN had incorporated (FNPEG) or FN had encapsulated (PEG+FN). **(e)** Water absorption of 3, 5 and 10 wt.% PEG only and FNPEG hydrogels (mean ±SD, n = 4). **(f)** Elastic modulus measured by AFM nanoindentation of 3, 5 and 10 wt.% PEG only and FNPEG hydrogels (mean ±SD, n > 100 curves). **(g)** FNPEG hydrogels with different ratios of degradable crosslinker (VPM peptide) (mean ±SD, n > 100 curves, *p-value<0.05, ANOVA test followed by a Tukey’s post hoc test). **(h)** Degradation profile of FNPEG with different ratios of degradable crosslinker (mean ±SD, n = 3). FN was covalently incorporated into PEG hydrogels, which could be further tuned to control their stiffness and degradability.

Prior to hydrogel formation, the disulphide bonds that hold the FN dimers together were reduced to obtain two FN monomers. FN monomers were then PEGylated via a Michael-type addition reaction and via functionalisation of the FN cysteine residues with controlled densities of 4-arm-PEG-maleimide molecules. On testing the biological activity of the PEGylated FN (Figure S1), we found it had similar availability of the cell adhesion-binding site and the heparin II-binding domain (Figure S1c, d) to native FN. As expected, differences between native and PEGylated FN were found mainly at the collagen binding site, which was less available in the PEGylated version (Figure S1e) relative to the native form. This is due to PEG molecules principally binding to the FN type I and II repetitions, which are the domains that contain cysteine residues (Figure 1a). We did not observe differences between the native and PEGylated FN forms when assessing adhesion and focal adhesion formation in C2C12 cells, demonstrating that these cells can fully interact with the protein after its PEGylation (Figure S1f-m). These findings agree with previous work reporting that PEGylated FN retains biological activity in terms of cell adhesion and FN fibril assembly, together with some proteolytic stability^56,57^. It is also noteworthy that full-length proteins are reported to retain high biological activity after the PEGylation process when incorporated into hydrogel systems^58–60^.

We then assessed the homogeneous distribution of FN within the hydrogel network via immunostaining (Figure 1c and Figure S2a). Hydrogels without FN did not show any staining. We also tested the efficiency of binding between FN and PEG by studying the release of FN from the hydrogels after 24 hours (Figure 1d and Figure S2b). Hydrogels into which PEGylated FN had incorporated (i.e., through covalent binding) did not release FN (FNPEG, Figure 1d, Figure S2b), whereas PEG hydrogels into which native FN was encapsulated (i.e., simply trapped) released after 24 hours (PEG+FN, Figure 1d).

Using PEG as a hydrogel network allows the physicochemical properties of the system to be controlled ^61–63^. Increasing the amount of PEG in the system (from 3 to 10 wt.%) increases water absorption in the hydrogels (Figure 1e) and increases Young’s modulus (Figure 1f), both independently of FN, the concentration of which was kept constant in the hydrogels at 1 mg/mL. These results agree with previously published results on standard PEG hydrogels, where an increase in the PEG percentage resulted in an increase in the Young’s modulus 53,61,64,65. PEG hydrogels can also be engineered to be cell-degradable^65^. To demonstrate controlled levels of degradation, we incorporated a protease degradable crosslinker (VPM peptide, Figure 1a) in combination with thiolated PEG. The degradation rate of hydrogels upon collagenase type I treatment increased with increasing amounts of VPM crosslinker, both for FN-containing hydrogels, FNPEG, and PEG hydrogels without FN (Figure 1h and Figure S2d). The addition of increasing ratios of the VPM peptide did not affect either the water sorption ability of the hydrogels (Figure S2c) nor their mechanical properties (Figure 1g).

Together, these data demonstrate that full-length and functional FN can be incorporated into a synthetic hydrogel system that has controlled stiffness and degradation rates.

### Full-length FN hydrogels sequester growth factors

As discussed, FN can promiscuously bind GFs, such as VEGF, through the heparin II-binding domain as long as this domain is available for interaction^13,20,22^. We demonstrate the ability of FN hydrogels to actively sequester GFs, compared to PEG only hydrogels (Figure 2). VEGF was fluorescently labelled to track its release over time (Figure 2a). FNPEG hydrogels retain up to 50% of the VEGF that was initially loaded into the hydrogel, compared to the 100% release of VEGF from PEG only hydrogels after 24h (Figure 2b). We also note that the presence of VPM peptide did not affect VEGF’s sequestration (Figure 2c), which demonstrates that FNPEG hydrogels can be engineered to control GF presentation independently of their degradability.

**Figure 2.**
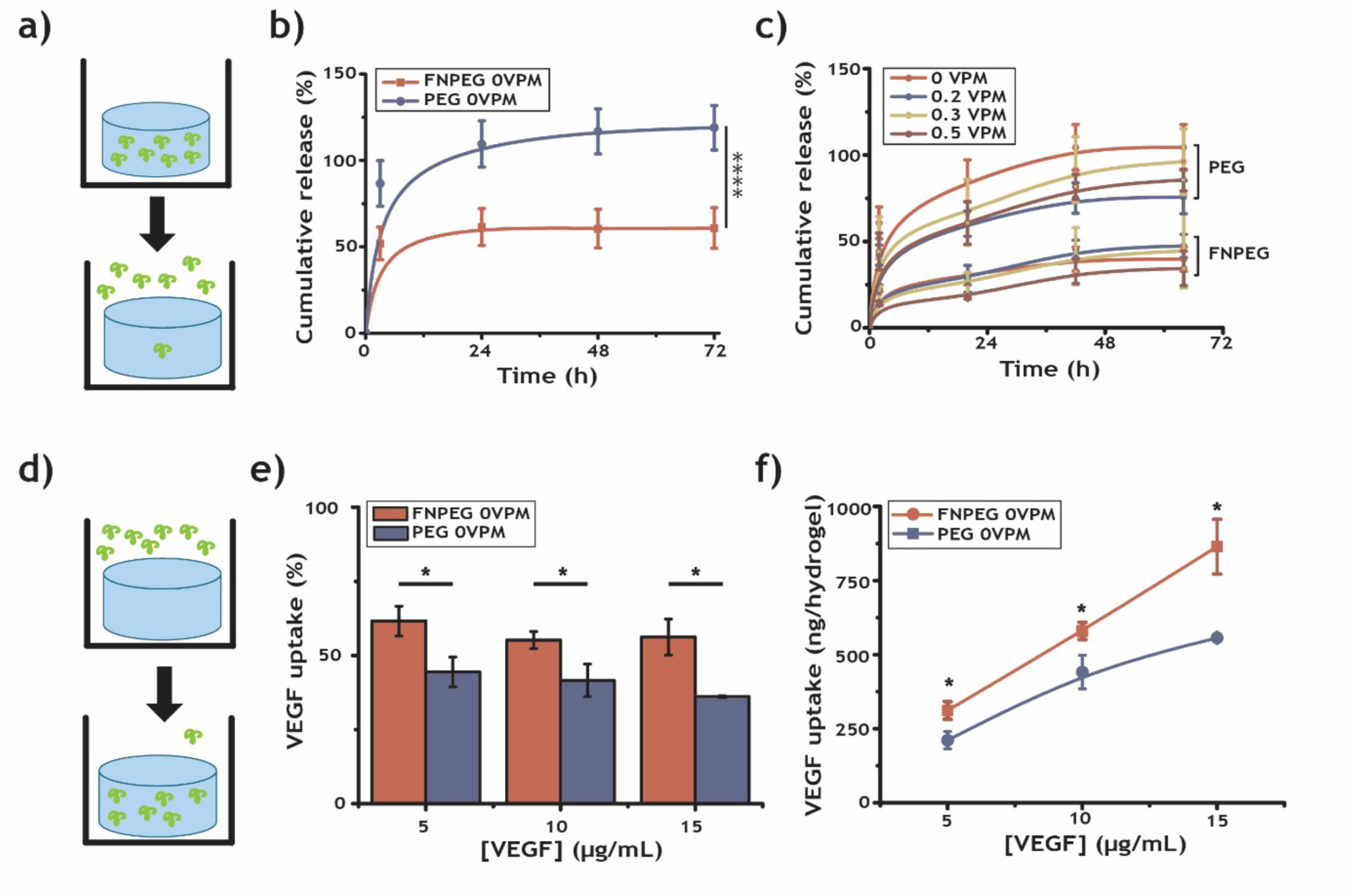
FN hydrogels actively bind VEGF. **(a)** Schematic of a release experiment in which the hydrogel is loaded with fluorescently labelled VEGF. The hydrogel was then immersed in DPBS and the fluorescent intensity of the supernatant tracked through time. **(b)** The cumulative release of fluorescently labelled VEGF from PEG only and FNPEG hydrogels, loaded with 10 µg/mL VEGF (mean ±SD, n = 6, ***p-value < 0.001, Kruskal-Wallis and Dunn’s post hoc test). **(c)** The cumulative release of fluorescently labelled VEGF from PEG only and FNPEG hydrogels with different ratios of degradable crosslinker (VPM) (mean ±SD, n = 3). **(d)** Schematic of the VEGF uptake assay, in which a hydrogel was immersed in fluorescently labelled VEGF solution and the fluorescence intensity of the supernatant measured. **(e)** Percentage of VEGF absorbed (*p-value < 0.05, Kruskal-Wallis with a Dunn’s post hoc test) and **(f)** amount of VEGF absorbed by PEG only and FNPEG hydrogels (*p-value < 0.05, Kruskal-Wallis and Dunn’s post hoc test). FNPEG hydrogels incorporate VEGF that is stably bound to FN in the 3D environment.

The capability of FNPEG hydrogels to sequester VEGF was further demonstrated in an uptake assay. In this assay, hydrogels were immersed in solutions that contained fluorescently labelled VEGF and their absorption was then measured (Figure 2d). FN hydrogels absorbed more VEGF compared to PEG only hydrogels for all concentrations of the initial VEGF solution used (Figure 2e, f). The increased uptake of VEGF in the FNPEG hydrogels cannot be explained by simple passive diffusion of the GF through the PEG network alone, since both PEG and FNPEG hydrogels absorbed similar amounts of water (Figure 1e). This demonstrates that FN binds VEGF as it diffuses from the initial solution, and that VEGF becomes part of the ‘solid phase’ of the hydrogel, and allows more GF to diffuse until equilibrium is reached. The difference in VEGF uptake between FNPEG and PEG, as shown in Figure 2f, for each concentration must be from the VEGF that is bound to FN in the hydrogel. Together, these data show the solid-phase presentation of GFs from full-length 3D FN crosslinked into a synthetic hydrogel with controlled stiffness and degradation rates. In the subsequent experiments, we used only 5 wt.% FNPEG and PEG gels with a ratio of 0.5 VPM.

### Full-length FN hydrogels promote microvasculature growth

We next assessed whether VEGF-loaded FN hydrogels promote endothelial cell activation and microvasculature growth. Human umbilical vein endothelial cells (HUVECs) were encapsulated, in situ, in both PEG only and FNPEG hydrogels, and retained high cell viability (Figure 3a), as expected, due to the mild crosslinking reaction of the hydrogels^53,54^. To investigate endothelial sprouting in 3D, we seeded HUVECs onto microcarrier beads. These cell-coated beads were then encapsulated within FNPEG hydrogels or in Matrigel, which was used as a positive control (Figure 3b). FNPEG hydrogels promoted high levels of sprouting, similar to those observed in Matrigel (Figure 3c). We note that for both FNPEG hydrogels and Matrigel sprouting only occurred in the presence of 500 ng/mL VEGF. The use of 50 ng/mL resulted in no sprouting in either FNPEG or Matrigel conditions.

**Figure 3.**
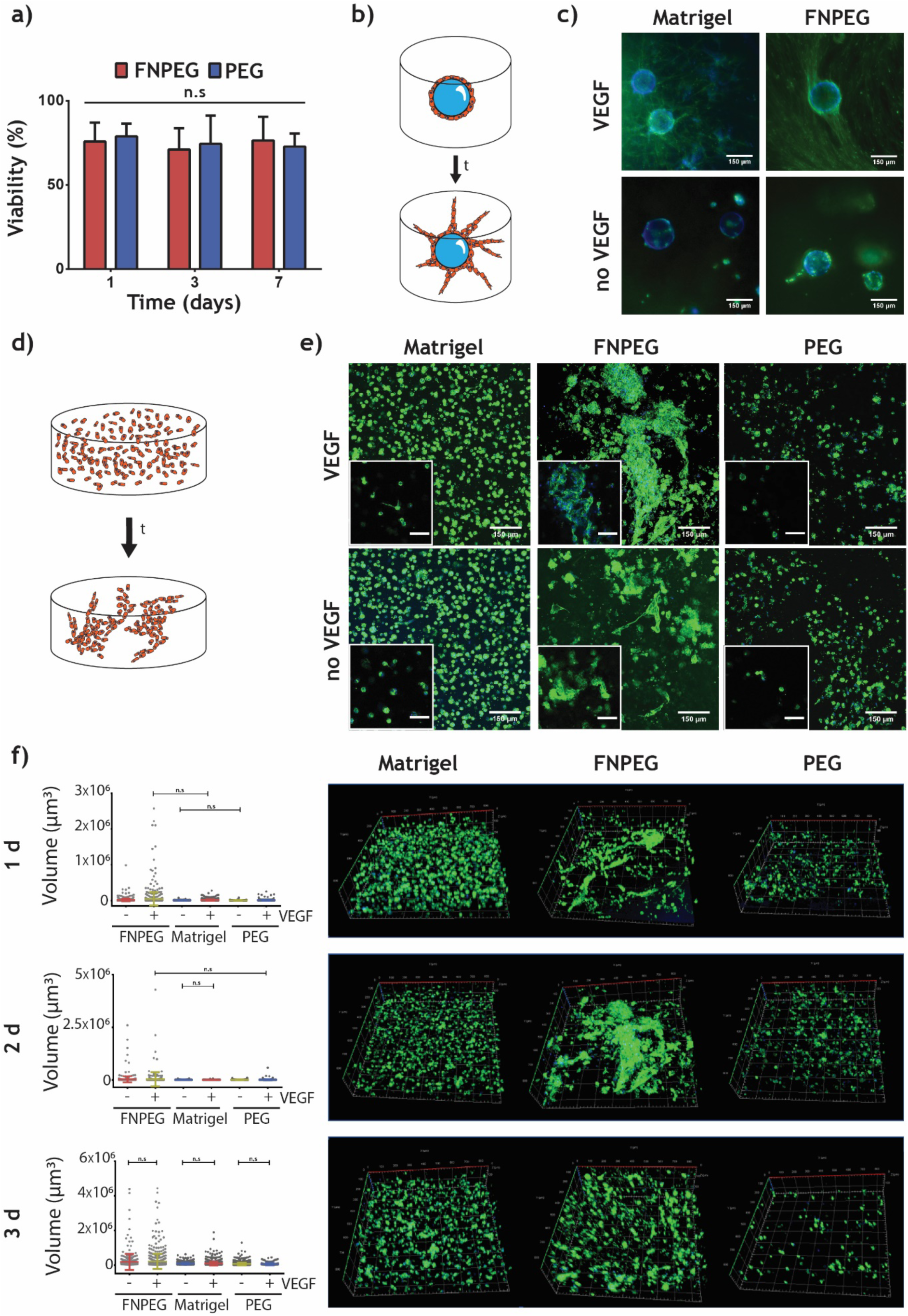
FN hydrogels promote sprouting and microvasculature growth in 3D. **(a)** Percentage viability of HUVECs after encapsulation in FNPEG and PEG only hydrogels. **(b)** Schematic of 3D sprouting assay, in which HUVECs were seeded onto collagen-coated dextran beads and the cell-coated beads encapsulated in FNPEG or Matrigel hydrogels. **(c)** Representative images of actin cytoskeleton (green) and nucleus (blue) of endothelial cell sprouting within FNPEG and Matrigel hydrogels (scale bar: 150 µm). **(d)** Schematic of endothelial cell 3D reorganisation assay, in which HUVECs were encapsulated within VEGF-loaded or no-VEGF FNPEG and PEG only hydrogels. **(e)** Representative z-axis projections of stacks at day 2 post-encapsulation (insets: representative slices of stack images at the same magnification, actin cytoskeleton (green and nucleus (blue)), scale bar: 150 µm. **(f)** (left) Volume of objects (µm^3^) at days 1, 2 and 3 after encapsulation, as quantified from stack images, and (right) representative 3D reconstructions of HUVECs within Matrigel, FNPEG or PEG at days 1, 2 and 3 (with VEGF) (mean ±SD, n > 100 objects). FN hydrogels are cytocompatible and promote endothelial cell sprouting in 3D at levels comparable to Matrigel.

Next, we investigated whether FNPEG gels promote the ability of HUVECs to activate and reorganise themselves in a 3D context, as happens in Matrigel (Figure 3d). To do so, we encapsulated HUVECs and VEGF within FNPEG hydrogels, as well as within Matrigel and PEG controls, and tracked cell morphology at early timepoints. We found that endothelial cells underwent more extensive morphological and structural changes within the VEGF-containing FNPEG hydrogels, as compared to the cells in the VEGF-containing Matrigel and PEG only hydrogel (Figure 3e, f and Figure S3). At days 1 and 2 after encapsulation, endothelial cells within the VEGF-containing FNPEG hydrogels formed multicellular and interconnected structures, as demonstrated in the 3D reconstruction images and volume quantification (Figure 3f). Endothelial cells did not form these complexes in PEG only (with or without VEGF), although cells in VEGF-containing Matrigel did show some multicellular structures, consisting of 3-4 cells at days 2 and 3. At day 3, the endothelial cell clusters formed in VEGF-containing FNPEG hydrogels started to disassemble, suggesting that these cellular complexes were not stable. During vasculogenesis, endothelial cells are highly dynamic and come together to form primitive capillary structures^66^. However, they require guidance from other cell types, such as pericytes and smooth muscle cells, to stabilise the newly formed capillary^67^. These experiments demonstrate that FNPEG hydrogels sequester VEGF and induce endothelial cells to form the early structures associated with vascularisation, more extensively than in cells cultured in Matrigel.

After showing that FNPEG hydrogels promote endothelial cell sprouting in 3D, we tested the system in a more complex environment using the chick chorioallantoic membrane (CAM) assay (Figure 4). Hydrogels (FNPEG hydrogel and PEG control) loaded with VEGF (and controls without VEGF) were placed on top of the exposed CAM, and chick embryos were incubated for four days. Images of the CAM were then taken and quantified. The results show that increased capillary formation occurred when FNPEG (with or without VEGF) was present, as can be seen by the number of branches and junctions counted per field (Figure 4c, d). CAMs incubated with VEGF-containing PEG only lacked extensive capillary formation, as seen in the empty condition (when a CAM was exposed but no material was placed on top). This is likely to be due to a rapid release of VEGF from PEG during the first hours, causing VEGF to be lost from the local environment relatively quickly. This would be in accordance with our VEGF release studies (Figure 2b, c), which showed that VEGF is rapidly released from PEG only hydrogels during the first 24 h. In contrast, FNPEG hydrogels (with or without VEGF) showed the highest levels of capillary formation. This is likely to be due to FNPEG hydrogels providing a more sustainable release of VEGF (as shown in our release studies, Figure 2b, c) but also because of the likely sequestration of natural VEGF present in the developing CAM ^68–70^, as also demonstrated in our in vitro VEGF uptake tests (Figure 3e, f).

**Figure 4.**
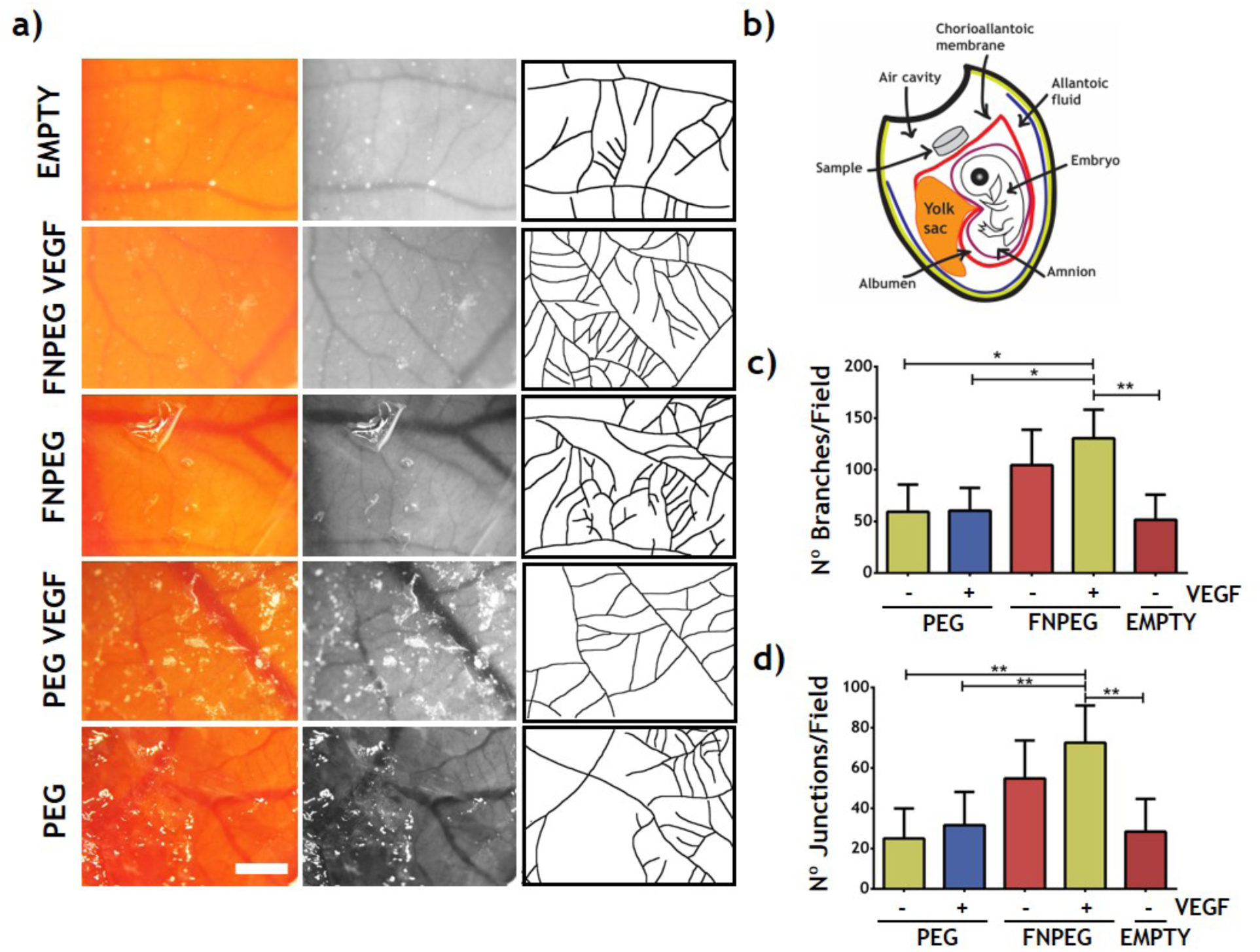
FN hydrogels promote capillary formation in vivo. **(a)** Representative images of chick chorioallantoic membrane (CAMs) (left column), green microscopy channel used for quantification (middle column), and (right column) manually drawn images of capillary branching. Conditions tested from top to bottom: “empty” (no hydrogel), “FNPEG VEGF” (FNPEG 0.5 VPM loaded with 125 ng VEGF), “FNPEG” (FNPEG 0.5 VPM), “PEG VEGF” (PEG only 0.5 VPM loaded with 125ng VEGF) and “PEG” (PEG only 0.5 VPM) (scale bar: 1 mm). **(b)** Schematic of the chick CAM assay, in which the egg shell is opened to expose the CAM and hydrogel samples are carefully placed on top. **(c)** Number of branches per image (mean ±SD, n > 10 images) and **(d)** number of junctions per image (mean ±SD, n > 10 images). FN hydrogels promote capillary formation in the chick chorioallantoic membrane assay.

### BMP2-loaded FN hydrogels promote bone formation

We used a non-healing (critical size), radial bone defect model in the adult mouse to demonstrate that FNPEG hydrogels promote bone formation in vivo when loaded with low concentrations of BMP2 (Figure 5). In this model, a 2.5 mm defect was introduced into the radial bone that does not heal spontaneously. The ulna was left intact to provide mechanical stabilisation and to avoid the use of additional external fixation plates. Hydrogel-filled implant tubes, 4 mm in length and that contain multiple pores along their sides^21,22,71^, were placed inside the bone defect.

**Figure 5.**
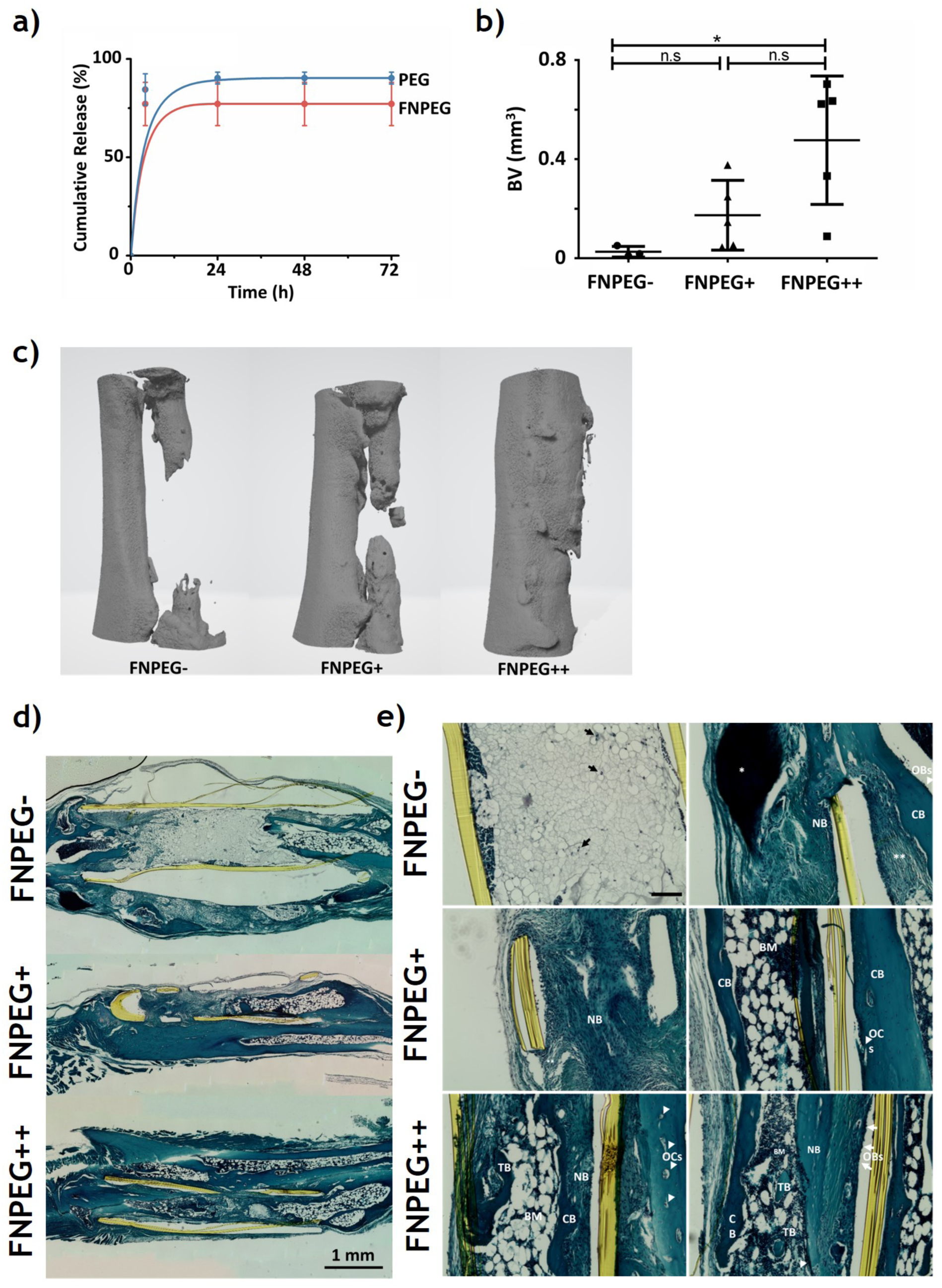
FN hydrogels promote bone growth in vivo. **(a)** Cumulative release of fluorescently labelled BMP2 from FNPEG hydrogels (%, mean ±SD, n = 4). **(b)** The percentage of bone volume (BV) quantified in FNPEG-(no BMP2, control), and in FNPEG+ and FNPEG++ hydrogels, which were initially loaded with 4.5 and 67.5 ng of BMP2, respectively (mean ±SD, n = 5, *p-value 0.0086). **(c)** 3D reconstructions of micro computed tomography (µCT) scans showing 4 mm of the ulna (left) and the radius (right), where the implant tube was placed. **(d)** Whole mount, longitudinal sections of representative forearms, with the proximal end to the left (scale bar: 1 mm). Yellow colour shows the walls of the implant tube. **(e)** Representative histological images of bone repair in each treatment group and the FNPEG-control (showing safranin O, fast green and haematoxylin staining) (scale bar: 100 µm). Black arrows show cell infiltration, * indicates bone deposition, ** marks fibrotic tissue, NB shows new bone formation, CB shows compact bone, OBs are osteoblasts indicated by white arrows, OCs are osteoclasts indicated by white arrowheads, TB shows trabecular bone, and BM shows bone marrow cavities. These results show that FN hydrogels promote bone growth at low BMP2 concentrations.

To use this model, we first studied the release of BMP2 from the hydrogel system in vitro (Figure 5a). FNPEG hydrogels released approximately 70% of the BMP2 initially loaded into them in the first 4 h, whereas PEG only hydrogels (that do not contain FN) released more than 90% of the BMP2 loaded into them in the same time period. We hypothesised that the other fraction of BMP2 in FNPEG hydrogels remained bound to FN.

We used two initial concentrations of BMP2 for this experiment: 5 and 75 µg/mL (denoted as FNPEG+ and FNPEG++, respectively). Implant tubes were filled with FNPEG hydrogels loaded with BMP2 the day before the experiment (hydrogels without BMP2 were used as a control, denoted FNPEG-). Taking into account that 70% of BMP2 (not bound to FN) is released during the first hours, the amount of BMP2 that remained within the FNPEG+ hydrogels was only 4.5 ng per implant tube for an initial loading of 5 µg/mL (providing an actual BMP2 concentration of 1.5 µg/mL retained within the hydrogel), and for FNPEG++ hydrogels, 67.5 ng of BMP2 per implant tube for an initial loading of 75 µg/mL (providing an actual BMP2 concentration of retained 22.5 µg/mL within the hydrogel). We note that 4.5 ng of BMP2 in this model is considered to be an ultralow dose^21,22^.

An analysis of the bone defects implanted with FNPEG++ hydrogels, by micro computed tomography (µCT) scans, showed that the defect was bridged in 2 out of 5 mice tested. When bone defects were implanted with the FNPEG+ hydrogels, varying results were recorded, and no closure of the bone gap was observed (Figure 5c). Likewise, the bone gap remained unclosed in the FNPEG-control (Figure 5c). Quantification of bone volume (BV, mm^3^) showed that the highest increase in bone formation occurred in the FNPEG++ implanted hydrogels, relative to the FNPEG-(no BMP2) controls (Figure 5b). Some FNPEG++-implanted bones also showed fusion to the ulna (Figure 5c). This could be due to the hydrogel swelling and coming into contact with the ulna through the pores of the implant tube. Some bone growth was also observed coming from the ulna in the FNPEG+ condition. Although no bone-defect closure was observed in the FNPEG+ group, we observed a trend of increased bone growth from FNPEG-, to FNPEG+ and to FNPEG++ that corresponded with increasing amounts of BMP2 in the implants (Figure 5b).

Longitudinal sections of the paraffin embedded forearms were stained with safranin-O, fast green and haematoxylin (Figure 5d, e). Figure 5d shows a whole mount of the implant tube position and greater collagen deposition (stained in blue) in the FNPEG+ and to FNPEG++ implanted bones, relative to the FNPEG-control, which did not stain for collagen around the defect. Figure 5e shows the formation of new tissue within the implant tube in more detail. Cell infiltration (as denoted by black arrows in Fig 5e) was observed in the FNPEG- hydrogels along the defect. The structures seen in this condition appear to resemble fat tissue. Some bone matrix deposits were observed at the outer part of the implant, probably to mechanically support the area. This is supported by the presence of osteoclasts at the proximal end of the defect (Figure S4), which would increase bone resorption in that area (Figure 5d).

In the FNPEG+ condition, compact bone formation was observed along the wall of the implant tube, with osteoblasts present at the outer layer and bone resorption was also observed, as indicated by the presence of osteoclasts (Figure 5e). This indicates that bone remodelling occurred within the implant tube. Bone marrow cavities were also present, which could indicate normal bone structure formation (Figure 5e).

In the FNPEG++ condition, implant tubes were completely filled with compact bone, as well as with new bone and trabecular bone, which filled the gap (Figure 5d, 5e). The presence of newly formed bone, together with that of bone marrow cavity structures, demonstrates that normal bone growth occurred within the implant tube. An endosteum-like structure was observed next to the implant tube wall, which was surrounded by osteoblasts, osteoclasts and compact bone. These results suggest that low doses of BMP2 within these matrices can support normal bone growth.

## Discussion

We have developed a platform to engineer hydrogels that incorporate full-length FN within the polymer network and that display controlled physico-chemical properties. The FN hydrogel system reported here is synthetic and tuneable, as shown by the characterisation of its mechanical properties and degradation profiles. We demonstrate that full-length FN can be covalently linked to these hydrogels without altering their key biological activities. We also demonstrate that FN hydrogels sequester VEGF from the environment and promote vascularisation efficiently, both in vitro and in vivo. FN hydrogels, when loaded with BMP2, can also promote bone growth in a non-healing mouse bone defect model, even at low BMP2 concentrations.

It remains for future studies to address whether bone regeneration could be improved by further optimising the initial amount of BMP2 loaded into the implant tube, and whether the generation of stiffer hydrogels could promote osteogenesis by the stem cells available at the fracture site.

Overall, our findings demonstrate that growth factors can be efficiently presented in a highly controllable, fully synthetic, 3D microenvironment for use in multiple tissue-engineering applications. The FN hydrogel system has the potential to substitute Matrigel in 3D culture models, given that it is a chemically defined system that contains full-length proteins and distinct, solid-phase, presentation of growth factors. This system can also be produced using purified human recombinant proteins, removing the need to screen for pathogens as occurs when using Matrigel matrices. The chemistries used to form the hydrogels are mild, cytocompatible and spontaneous at physiological pH and temperature. Moreover, the created hydrogels are transparent, and so are well-suited for colorimetric/fluorometric assays. Furthermore, being a biosynthetic system, it has the potential to be more amenable for translation in e.g. drug testing platforms or tissue engineering ^72^.

## Methods

### Fibronectin PEGylation

Fibronectin (FN, YoProteins, 3 mg/mL) was PEGylated by modifying a procedure from Almany et al. ^58^.

### Hydrogel Formation

PEG hydrogels were formed using Michael-type addition reaction under physiological pH and temperature following protocol from Phelps et al.^53^. Briefly, a final concentration of 1 mg/mL of PEGylated FN was added to different amounts of PEGMAL (3 wt.%, 5 wt.% or 10 wt.%). The thiolated crosslinker was added always at the end, at a molar ratio 1:1 maleimide:thiol to ensure full crosslinking. The crosslinkers used were either PEG-dithiol (PEGSH, 2 KDa, Creative PEGWorks) or mixtures of PEGSH and protease-degradable peptide, flanked by two cysteine residues (VPM peptide, GCRD<u>VPM</u>SMRGGDRCG, purity 96.9%, Mw 1696.96 Da, GenScript). Cells and/or soluble molecules, such as GFs, were always mixed with the protein and PEGMAL before addition of the crosslinker. Once the crosslinker was added, samples were incubated for 30 min at 37°C to allow gelation. PEG-only hydrogels were produced as well without the addition of the PEGylated FN. The nomenclature used in this manuscript is: x% (FN)PEG yVPM, x being the percentage of PEGMAL used and y the ration of VPM peptide crosslinker added. When not indicated, hydrogels were 5% FNPEG 0.5VPM.

### Fibronectin immunostaining

Hydrogels (3, 5 and 10 wt.% FNPEG or PEG only as a control) were embedded in optimal cutting temperature compound (O.C.T compound, VWR) and flash frozen by immersion in liquid nitrogen to preserve the structure of the gel. Samples were stored at −80°C until use. A cryostat (Leica, −20°C) was used to cut the samples, and sections 100 µm in thickness were prepared on treated microscope slides (ThermoFisher).

FN was detected via immunofluorescence in hydrogel cryosections. Sections were blocked with blocking buffer (1% Bovine Serum Albumin, BSA, Sigma) for 30 min at RT. Then, primary antibody rabbit polyclonal-anti-FN (Sigma, 1:200) was added and incubated for 1 h at RT. Samples were washed three times using washing buffer (0.5% Tween 20 (Sigma) in PBS). Then, secondary antibody goat-anti-rabbit-Cy3 (Jackson ImmunoResearch, 1:400) was incubated for 1 h at RT and protected from light. Samples were washed three times using washing buffer and were mounted with VECTASHIELD mounting media without DAPI (Vector Laboratories). Images were taken with a ZEISS AxioObserver Z.1 at different magnifications.

### Fibronectin release

Hydrogels (either 5% FNPEG, 5% PEG or 5% PEG hydrogels with native FN entrapped) were immersed for 24 h in PBS to assess the release of FN. Solutions from supernatants were collected and quantified using bicinchoininc acid colorimetric assay (MicroBCA assay kit, ThermoFisher Scientific),following the manufacturer’s instructions. Briefly, BSA standards and samples were loaded onto a 96-well microplate and were mixed with the working reagent. Then, the microplate was sealed and incubated for 2 h at 37°C. After incubation, the microplate was let to cool down for 20 min at RT, protected from light. The absorbance at 562 nm was measured using a plate reader (BIOTEK). Conditions were prepared in triplicate and samples were measured in triplicate.

### Water absorption

Hydrogels were formed and immersed in milliQ water for up to a week using filtered tubes. For every timepoint, samples were centrifuged at 6500 rpm for 5 min to remove the supernatant from the basket so that only the hydrogel sample remained in the basket holder. Then, sample and basket were weighed, and the mass of the empty basket was subtracted. The amount of water absorbed was calculated as follows:

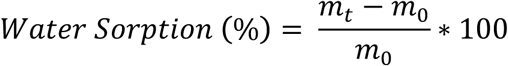

Where *m*_*t*_ is the weight of the hydrogel at a certain time and *m*_*0*_ the weight of the hydrogel after formation.

### Mechanical properties

Nanoindentation was assessed using atomic force microscopy in force spectroscopy mode (AFM/FS, Nanowizard-3, JPK). Cantilevers (Arrow-TL1-50, spring constant ∼ 0.03 N/m, Nano World innovative technologies) were functionalised manually with silicon oxide microbeads (20 mg/mL, 20 µm diameter, monodisperse, Corpuscular Inc.). The actual stiffness of the cantilever was estimated using the thermal calibration method. Samples tested were 100 µm cryosections fully swollen in milliQ water. Measurements were carried out in immersion. Indentation of at least 500 nm were assessed using constant force. The area of the sample was mapped defining squared areas (2500 µm^2^, 25 measurements). Five maps per replicate were measured and at least three replicates per sample were tested, unless otherwise noted. The analysis (JPK-SPM processing software) was performed using the Hertz model for a spherical indenter to fit the curves obtained.

### Degradation assays

Hydrogels were formed and swollen overnight in PBS. All samples were weighed before starting the degradation. Then, samples were covered with protease solution (collagenase type I, Gibco, 50 U/mL in PBS, 37°C). At each timepoint, all supernatant was removed by centrifugation at 6500 rpm for 5 min and samples were weighed. To continue the experiment, fresh protease solution was added. The degradation rate was calculated as follows:

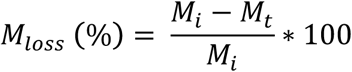

Where M_loss_ is the percentage of mass lost during degradation, M_i_ the mass after swelling in milliQ water (initial mass), and M_t_, the mass at the different timepoints after the addition of the protease solution.

### Growth factor labelling

In order to study GF binding and release, VEGF or BMP2 (carrier free, R&D Systems) were fluorescently labelled with an amino reactive dye (DyLight® NHS Ester, Thermo Fisher Scientific) following the manufacturer’s instructions. Briefly, GFs were dialysed (Mini-A-Lyzer, COMW 10 kDa, ThermoFisher) against 0.05 M Sodium borate buffer at pH 8.5 for 2 h at RT. Then, the appropriate amount of dye was added (as calculated by the manufacturer’s guidelines) to the GF solution. The dye and the GF were let to react for 1 h at RT, protected from light. Then, the non-reacted dye was removed by dialysis against PBS for three hours. The labelled VEGF was aliquoted and stored at −20°C until use. Fluorescently labelled VEGF or BMP2 were named VEGF-488 or BMP2-488, respectively, in the main text.

### Growth factor release

Hydrogels were prepared, as described in hydrogel formation section above, that incorporate VEGF-488 or BMP2-488. The final concentration of labelled GF loaded was 10 µg/mL. Then, samples were immersed in PBS and incubated at 37°C protected from light. At each timepoint, all the PBS solution was taken and used to measure the fluorescence (Ex/Em 493/518 nm) using a plate reader (BIOTEK). Fresh PBS was added after each timepoint. A standard curve using VEGF-488 or BMP2-488 was prepared and measured together with the samples. An empty condition (not loaded with GF) was used as control. All conditions were prepared in triplicate, and each sample was measured three times. The cumulative release (%) was calculated as follows:

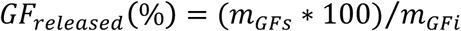

Where GF_released_ is the percentage of VEGF-488 or BMP2-488 released from hydrogels, m_GFs_ is the amount of VEGF-488 or BMP2-488 measured in the supernatant and m_GFi_ is the amount of VEGF-488 or BMP2-488 initially loaded.

### VEGF uptake

Hydrogels were immerse in 10 mM L-cysteine solution for 2 h to ensure that all the maleimide groups from the hydrogels were reacted. After that, samples were washed three times in PBS and immersed in VEGF-488 solutions of different concentrations (5, 10 and 15 µg/mL). Once immersed, samples were incubated for 20 h at 37°C while protected from light. The supernatant was then taken and read using a plate reader (Ex/Em 493/518 nm). All conditions were prepared in quadruplicate and all samples were measured twice. The initial solutions were also measured and used as standard curve to be able to correlate fluorescence intensity with VEGF-488concentration. The fluorescence intensity of hydrogels immersed in PBS (without VEGF-488) was measured to normalise the data. The percentage of VEGF-488 absorbed was calculated as follows:

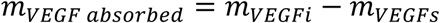

Where m_VEGF absorbed_ is the final mass of VEGF-488 retained in the hydrogels, m_VEGFi_ is the initial total mass of VEGF-488 added to the samples in solution, and m_VEGFs_ is the total mass of VEGF-488 measured in the supernatant after incubation. Once the amount of VEGF-488 retained in the samples was calculated, the percentage of VEGF-488 absorbed was calculated as follows:

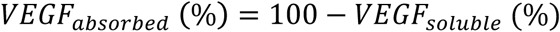

Where VEGF_absorbed_ is the percentage of VEGF retained in the hydrogel and VEGF_soluble_ is the percentage of VEGF measured from the supernatants after the incubation of VEGF with the hydrogel.

### Cell culture

Human umbilical vein endothelial cells (HUVECs) (Caltag Medsystems, passage 1-5) were used for viability and angiogenesis/vascularisation *in vitro* assays. HUVECs were grown in growth media (large vessel endothelial cell (LVEC) medium, Caltag Medsystems). Human dermal fibroblasts (HDFs) (Caltag Medsystems, passage 1-6) were used for angiogenesis studies. Both cell types were grown in Dulbecco’s modified Eagle’s medium (DMEM, Gibco) high glucose without pyruvate and 10% fetal bovine serum (FBS, Gibco) until seeding. Once seeded, LVEC media was used. All media used were supplemented with 1% penicillin/streptomycin (Gibco).

### Cell viability

Cytocompatibility of hydrogels was tested using the Live/Dead assay (ThermoFisher), according to the manufacturer’s instructions. Briefly, HUVECs cells were encapsulated at 10^6^ cells/mL to allow single-cell analysis and were incubated within the hydrogels at different timepoints. For each timepoint, cells were stained with 2 µM Calcein-AM and NucBlue and incubated for 15 min. Samples were washed twice before imaging with a confocal microscope (ZEISS CLSM 880) at 10X magnification.

### Angiogenesis assays

Angiogenic sprouting was assessed using bead microcarriers^48^. Briefly, HUVECs were mixed with dextran-coated Cytodex 3 microcarriers (Sigma) at a final concentration of 400 cells per bead in one millilitre of growth medium. Cells and beads were mixed gently every 20 min for 4 h at 37°C. Then, cell-coated beads were transferred to a flask with growth medium and incubated overnight (37°C and 5% CO2). Before encapsulation, cell-coated beads were washed three times with growth medium. Finally, cell-coated beads were loaded into hydrogels at a final concentration of 400 beads/mL. Once the hydrogels were prepared, twenty thousand human dermal fibroblasts were seeded on top. Growth media supplemented with different concentrations of VEGF (0, 50, 500 ng/mL, R&D Systems) was changed every other day. The assay was monitored every day for four days. Samples were fixed using 4% para-formaldehyde for 30 min at RT and stained for actin (AlexaFluor-488 Phalloidin dilution 1:300) and for the nucleus (NucBlue, LifeTechnologies) for 1 h. Samples were washed three times with 0.5% Tween20 in DPBS and mounted onto glass bottom petri dishes using VECTASHIELD mounting medium (VectorLabs). Samples were imaged using a ZEISS Axio Observer Z1 and were prepared in triplicate.

### Vascularisation assays

For vascularisation studies, HUVECs were encapsulated (5·10^6^ cells/mL) *in situ* within hydrogels loaded with 200 pmol/mL VEGF-165 (R&D Systems) or within non-loaded hydrogels (without GF). Samples at days one, two and three were fixed with 4% paraformaldehyde for 30 min at RT and stained for actin and the nucleus. Samples were imaged using confocal microscopy (ZEISS LSM 880) at 10X magnification. Stacks obtained from confocal imaging were analysed using ImageJ 1.51v. Briefly, actin cytoskeleton stacks were opened and segmented using the trainable Weka segmentation 3D plugin. Once the stacks were converted to 8-bit segmented stacks, the segmented objects were quantified using the 3D objects counter tool with the following parameters: volume (V, µm3), number of voxels/object, surface (S, µm2), number of voxels/surface and centroids. Prior to the quantification, a size exclusion filter was applied, so objects smaller than 500 voxels were not counted (to avoid quantification of segmented background noise). The sphericity (Ψ) of the objects was calculated as follows: Ψ=([π^(1/3)·(6·V)^(2/3)])/S.

### Chick chorioallantoic membrane assay

Fertilised chick eggs were received at day seven post-fertilisation (E7). Eggs were kept in an incubator (37.5 °C, 50-60% relative humidity). To perform the chorioallantoic membrane (CAM) assay, eggs were candled to detect and mark the air sac of the chick embryo, and the egg shell was then cut on top of the air sac to expose the CAM. Once the membrane was exposed, each sample was laid carefully on top of the membrane. Samples tested were: 5% PEG 0.5VPM, 5%PEG 0.5VPM with 2.5 µg/mL VEGF165, 5% FNPEG 0.5VPM, 5% FNPEG 0.5VPM with 2.5 µg/mL VEGF165 and an empty condition, where the CAMs were exposed but no material was placed on top. After that, the exposed area of the egg with the sample was sealed and labelled. All eggs were incubated for four days (E12), when the membranes were imaged using a stereomicroscope (Leica MZ APO) using X8 and X16 magnification. Six replicates per condition were used; two pictures per replicate at each magnification used were taken. For the quantification, images at X16 magnification were used and the images were anonymised. The green channel of the RGB picture was chosen for the segmentation as it was the one with the best contrast to detect the capillaries. Segmentation was assessed manually, tracing a black line on top of each capillary. The result of the segmentation was used to quantify the number of branches, the number of junctions, and the number of triple points per image via the skeletonize tool on ImageJ.

### Murine non-healing bone defect model

All murine experiments were conducted under the Animals (Scientific Procedures) ACT 1986 (ASPel project license n° 70/8638) and all the research performed, complied with ethical regulations approved by the University of Glasgow’s ethical committee.

### Implant preparation

Polyimide implant tubes with lateral holes were used as sleeves to be filled with the hydrogel samples. FN hydrogels (5 wt.%, 0.5VPM) were loaded with BMP2 (R&D Systems) at a final concentration of 0, 5 or 75 µg/mL (FNPEG-, FNPEG+ or FNPEG++, respectively). The implants were prepared the day before surgery and were kept in PBS until use.

### Bone radial segmental defect surgery

This experiment was conducted under the Animals (Scientific Procedures) ACT 1986 (ASPel project license n° 70/8638). C57BL/6 male mice (8 weeks old, Charles River, n = 5 mice/condition) were anaesthetised using isoflurane gas. Under anaesthesia, mice were provided with a dose of buprenorphine and carprofen for pain relief. An incision on the skin was carried out along the forearm and the radius was exposed using a periosteal elevator. The centre of the radius was revealed in order to introduce a 2.5 mm wound in the bone, using a custom-made parallel double-bladed bone cutter. The ulna was left intact. The implant was then placed into the introduced bone defect, abutting its proximal and distal ends. Finally, the wound was closed with a degradable suture. Mice were monitored during the experiment for signs of distress, movement, and weight loss.

### Analysis of bone growth

Eight weeks after surgery, bone samples were explanted and fixed in 4% para-formaldehyde and immersed in 70% ethanol. Bone samples were analysed using microcomputer tomography (µCT, Bruker Skyscan Micro X-ray CT), and then decalcified using Krajian solution (citric acid, formic acid) for 3 d, until soft and pliable. They were then paraffin embedded for sectioning. Quantification of the bone volume was performed using the CTAn software (Bruker). In order to ensure that only new bone formation was measured, the volume of interest (VOI) was selected to evaluate a central 2.0 mm length of the 2.5 mm total defect size. Samples were paraffin embedded for histological analysis. Sections (of 7 µm thickness) were stained for haematoxylin-Safranin O-Fast Green. Briefly, Sections were deparaffinised and rehydrated in water. Then, Mayer’s haematoxylin staining was performed for 8 min and Scott’s solution was used for 1 min to blue up the nuclear staining. After that, samples were rinsed in 1% acid alcohol and water. Then, 0.5% fast green solution for 30 sec was used and sections were rinsed in 1% acetic acid for 3 sec. Finally, 0.1% safranin O solution was used for 5 min and sections were washed in 70%, 95% and absolute ethanol for 1 min each. Sections were cleared twice with Histo-Clear for 5 min and mounted with DPX mounting media. Sections were then imaged with an EVOS FL microscope at 20X magnification. Whole mounts were stitched together using the Image Composite Editor software.

### Statistical analysis

The statistical analysis was performed using GraphPad Prism 6.01 software. All in vitro experiments were carried out in triplicate unless otherwise noticed. All graphs represent mean ± standard deviation (SD) unless otherwise noted. The goodness of fit of all data-sets was assessed via D’Agostino-Pearson Normality test. When comparing three or more groups: normal distributed populations were analysed via analysis of variance test (ANOVA test) performing a Tukey’s post hoc test to correct for multiple comparisons; when populations were not normally distributed, a Kruskal-Wallis test was used with a Dunn’s post hoc test to correct for multiple comparisons. When comparing only two groups, parametric (normal distributed population, t-test) or non-parametric (Mann-Whitney test) tests were performed. Differences among groups are stated as follows: for p-values <0.05 (*), when p-values <0.01 (**), for p-values < 0.005 (***), for p-values < 0.001 (****), when differences between groups are not statistically significant (n.s).

## Acknowledgements

This study was supported by the UK Regenerative Medicine Platform (MRC grant MR/L022710/1), the UK Engineering and Physical Sciences Research Council (EPSRC EP/P001114/1) and a programme grant from Find A Better Way. μCT work was supported by the European Research Council (ERC) under the European Union’s Seventh Framework Programme (FP7/2007-2013) (grant agreement No. [615030]). S.T. acknowledges support from the University of Glasgow through their internal scholarship funding program.

## Author contributions

S.T., C.G-G., P.R-T., A.R. and J.W. performed the experiments; S.T. analysed the data; S.T., M. D. and M.S-S designed the experiments.; S.T., M.D. and M.S-S. wrote the paper.

## References

1. Dalby, M. J., García, A. J. & Salmeron-Sanchez, M. Receptor control in mesenchymal stem cell engineering. Nat. Rev. Mater. 3, (2018).

2. Hartikainen, J. Vascular Endothelial Growth Factors Biology and Current Status of Clinical Applications in Cardiovascular Medicine. 49, (2007).

3. Martino, M. M. et al. Extracellular Matrix and Growth Factor Engineering for Controlled Angiogenesis in Regenerative Medicine. Front. Bioeng. Biotechnol. 3, 1–8 (2015).

4. Mitchell, A. C., Briquez, P. S., Hubbell, J. A. & Cochran, J. R. Engineering growth factors for regenerative medicine applications. Acta Biomaterialia (2016). doi:10.1016/j.actbio.2015.11.007

5. Sprugel, K. H., McPherson, J. M., Clowes, A. W. & Ross, R. Effects of growth factors in vivo. I. Cell ingrowth into porous subcutaneous chambers. Am. J. Pathol. (1987).

6. Stejskalová, A., Oliva, N., England, F. J. & Almquist, B. D. Biologically Inspired, Cell-Selective Release of Aptamer-Trapped Growth Factors by Traction Forces. 1806380, (2019).

7. Crouzier, T., Ren, K., Nicolas, C., Roy, C. & Picart, C. Layer-by-layer films as a biomimetic reservoir for rhBMP-2 delivery: Controlled differentiation of myoblasts to osteoblasts. Small 5, 598–608 (2009).

8. Martino, M. M., Briquez, P. S., Maruyama, K. & Hubbell, J. A. Extracellular matrix-inspired growth factor delivery systems for bone regeneration. Adv. Drug Deliv. Rev. 94, 41–52 (2015).

9. Pike, D. B. et al. Heparin-regulated release of growth factors in vitro and angiogenic response in vivo to implanted hyaluronan hydrogels containing VEGF and bFGF. Biomaterials 27, 5242–5251 (2006).

10. Jha, A. K. et al. Matrix metalloproteinase-13 mediated degradation of hyaluronic acid-based matrices orchestrates stem cell engraftment through vascular integration. Biomaterials 89, 136–147 (2016).

11. Martino, M. M., Briquez, P. S., Ranga, A., Lutolf, M. P. & Hubbell, J. A. Heparin-binding domain of fibrin(ogen) binds growth factors and promotes tissue repair when incorporated within a synthetic matrix. Proc. Natl. Acad. Sci. 110, 4563–4568 (2013).

12. Laporte, L. De, Rice, J. J., Tortelli, F. & Hubbell, J. A. Tenascin C Promiscuously Binds Growth Factors via Its Fifth Fibronectin Type III-Like Domain. 8, (2013).

13. Martino, M. M. & Hubbell, J. A. The 12th-14th type III repeats of fibronectin function as a highly promiscuous growth factor-binding domain. FASEB J. 24, 4711–4721 (2010).

14. Wang, R. N. et al. Bone Morphogenetic Protein (BMP) signaling in development and human diseases. Genes Dis. 1, 87–105 (2014).

15. Martino, M. M. et al. Engineering the growth factor microenvironment with fibronectin domains to promote wound and bone tissue healing. Sci. Transl. Med. 3, (2011).

16. Wijelath, E. S. et al. Heparin-II domain of fibronectin is a vascular endothelial growth factor-binding domain: Enhancement of VEGF biological activity by a singular growth factor/matrix protein synergism. Circ. Res. 99, 853–860 (2006).

17. Wijelath, E. S. et al. Fibronectin promotes VEGF-induced CD34+cell differentiation into endothelial cells. J. Vasc. Surg. (2004). doi:10.1016/j.jvs.2003.10.042

18. Wijelath, E. S. et al. Novel vascular endothelial growth factor binding domains of fibronectin enhance vascular endothelial growth factor biological activity. Circ. Res. (2002). doi:10.1161/01.RES.0000026420.22406.79

19. Llopis-Hernández, V., Cantini, M., González-García, C. & Salmerón-Sánchez, M. Material-based strategies to engineer fibronectin matrices for regenerative medicine. Int. Mater. Rev. 60, 245–264 (2015).

20. Moulisová, V. et al. Engineered microenvironments for synergistic VEGF – Integrin signalling during vascularization. Biomaterials 126, 61–74 (2017).

21. Cheng, Z. A. et al. Nanoscale Coatings for Ultralow Dose BMP-2-Driven Regeneration of Critical-Sized Bone Defects. 1800361, (2019).

22. Llopis-hernández, V. et al. Material-driven fibronectin assembly for high-efficiency presentation of growth factors. 1–10 (2016).

23. Klass, C. M., Couchman, J. R. & Woods, A. Control of extracellular matrix assembly by syndecan-2 proteoglycan. J. Cell Sci. (2000).

24. Woods, A., Longley, R. L., Tumova, S. & Couchman, J. R. Syndecan-4 binding to the high affinity heparin-binding domain of fibronectin drives focal adhesion formation in fibroblasts. Arch. Biochem. Biophys. (2000). doi:10.1006/abbi.1999.1607

25. Guan, J. L. & Hynes, R. O. Lymphoid cells recognize an alternatively spliced segment of fibronectin via the integrin receptor α4β1. Cell (1990). doi:10.1016/0092-8674(90)90715-Q

26. Wayner, E. A., Garcia-Pardo, A., Humphries, M. J., McDonald, J. A. & Carter, W. G. Identification and characterization of the T lymphocyte adhesion receptor for an alternative cell attachment domain (CS-1) in plasma fibronectin. J. Cell Biol. (1989). doi:10.1083/jcb.109.3.1321

27. Hielscher, A., Ellis, K., Qiu, C., Porterfield, J. & Gerecht, S. Fibronectin deposition participates in extracellular matrix assembly and vascular morphogenesis. PLoS One 11, 1–27 (2016).

28. Zhou, X. et al. Fibronectin fibrillogenesis regulates three-dimensional neovessel formation. Genes Dev. (2008). doi:10.1101/gad.1643308

29. Pankov, R. Fibronectin at a glance. J. Cell Sci. 115, 3861–3863 (2002).

30. Magnusson, M. K. & Mosher, D. F. Fibronectin: Structure, assembly, and cardiovascular implications. Arterioscler. Thromb. Vasc. Biol. 18, 1363–1370 (1998).

31. Leiss, M., Beckmann, K., Girós, A., Costell, M. & Fässler, R. The role of integrin binding sites in fibronectin matrix assembly in vivo. Curr. Opin. Cell Biol. 20, 502–507 (2008).

32. Krishnakumar, G. S. & Roffi, A. Clinical application of bone morphogenetic proteins for bone healing : a systematic review. 1073–1083 (2017). doi:10.1007/s00264-017-3471-9

33. Cahill, K. S., Chi, J. H., Day, A. & Claus, E. B. Prevalence, complications and Hospital Charges Associated With Use of Bone-Morphogenetic Proteins in Spinal Fusion Procedures. J. Am. Med. Assoc. 302, 58–66 (2009).

34. James, A. W. et al. A Review of the Clinical Side Effects of Bone Morphogenetic Protein-2. Tissue Eng. Part B Rev. 22, 284–297 (2016).

35. Regranex. Available at: https://regranex.com/.

36. Shegarfi, H. & Reikeras, O. Review Article: Bone Transplantation and Immune Response. J. Orthop. Surg. (2016). doi:10.1177/230949900901700218

37. Myeroff, C. & Archdeacon, M. Autogenous bone graft: Donor sites and techniques. Journal of Bone and Joint Surgery - Series A (2011). doi:10.2106/JBJS.J.01513

38. Dimitriou, R., Mataliotakis, G. I., Angoules, A. G., Kanakaris, N. K. & Giannoudis, P. V. Complications following autologous bone graft harvesting from the iliac crest and using the RIA: A systematic review. Injury (2011). doi:10.1016/j.injury.2011.06.015

39. Hankenson, K. D., Dishowitz, M., Gray, C. & Schenker, M. Angiogenesis in bone regeneration. Injury 42, 556–561 (2011).

40. Gerber, H. P. & Ferrara, N. Angiogenesis and bone growth. Trends Cardiovasc. Med. 10, 223–228 (2000).

41. Kanczler, J. M. & Oreffo, R. O. C. Osteogenesis and Angiogenesis: the Potential for Engineering Bone. Eur. Cells Mater. 15, 100–114 (2008).

42. Almubarak, S. et al. Tissue engineering strategies for promoting vascularized bone regeneration. 83, 197–209 (2016).

43. Mittwede, P. N., Gottardi, R., Alexander, P. G., Tarkin, I. S. & Tuan, R. S. Clinical Applications of Bone Tissue Engineering in Orthopedic Trauma. 99–108 (2018).

44. Kolbe, M. et al. Paracrine Effects Influenced by Cell Culture Medium and Consequences on Microvessel-Like Structures in Cocultures of Mesenchymal Stem Cells and Outgrowth Endothelial Cells. Tissue Eng. Part A (2011). doi:10.1089/ten.tea.2010.0474

45. Wang, L. et al. Osteogenesis and angiogenesis of tissue-engineered bone constructed by prevascularized β-tricalcium phosphate scaffold and mesenchymal stem cells. Biomaterials (2010). doi:10.1016/j.biomaterials.2010.08.036

46. Ferrara, N., Gerber, H. P. & LeCouter, J. The biology of VEGF and its receptors. Nat. Med. 9, 669–676 (2003).

47. Leslie-Barbick, J. E., Moon, J. J. & West, J. L. Covalently-immobilized vascular endothelial growth factor promotes endothelial cell tubulogenesis in poly(ethylene glycol) diacrylate hydrogels. J. Biomater. Sci. Polym. Ed. 20, 1763–1779 (2009).

48. Nakatsu, M. N. et al. Angiogenic sprouting and capillary lumen formation modeled by human umbilical vein endothelial cells (HUVEC) in fibrin gels: The role of fibroblasts and Angiopoietin-1. Microvasc. Res. 66, 102–112 (2003).

49. E. Saik, J., K. McHale, M. & L. West J. Biofunctional Materials for Directing Vascular Development. Curr. Vasc. Pharmacol. 10, 331–341 (2012).

50. Leslie-barbick, J. E., Moon, J. J. & West, J. L. Covalently-Immobilized Vascular Endothelial Growth Factor Promotes Endothelial Cell Tubulogenesis in Poly (ethylene glycol) Diacrylate Hydrogels. 20, 1763–1779 (2009).

51. Phelps, E. A., Landazuri, N., Thule, P. M., Taylor, W. R. & Garcia, A. J. Bioartificial matrices for therapeutic vascularization. Proc. Natl. Acad. Sci. 107, 3323–3328 (2010).

52. Briquez, P. S., Clegg, L. E., Martino, M. M., Gabhann, F. Mac & Hubbell, J. A. Design principles for therapeutic angiogenic materials. Nature Reviews Materials 1, (2016).

53. Phelps, E. A. et al. Maleimide cross-linked bioactive PEG hydrogel exhibits improved reaction kinetics and cross-linking for cell encapsulation and in situ delivery. Adv. Mater. 24, 64–70 (2012).

54. Phelps, E. A., Templeman, K. L., Thulé, P. M. & García, A. J. Engineered VEGF-releasing PEG– MAL hydrogel for pancreatic islet vascularization. Drug Delivery and Translational Research 5, 125–136 (2015).

55. Jansen, L. E., Negrón-Piñeiro, L. J., Galarza, S. & Peyton, S. R. Control of thiol-maleimide reaction kinetics in PEG hydrogel networks. Acta Biomater. 1133, (2018).

56. Zhang, C., Ramanathan, A. & Karuri, N. W. Proteolytically stabilizing fibronectin without compromising cell and gelatin binding activity. Biotechnol. Prog. 31, 277–288 (2015).

57. Zhang, C., Desai, R., Perez-Luna, V. & Karuri, N. PEGylation of lysine residues improves the proteolytic stability of fibronectin while retaining biological activity. Biotechnol. J. 9, 1033–1043 (2014).

58. Almany, L. & Seliktar, D. Biosynthetic hydrogel scaffolds made from fibrinogen and polyethylene glycol for 3D cell cultures. Biomaterials 26, 2467–2477 (2005).

59. Francisco, A. T. et al. Photocrosslinkable laminin-functionalized polyethylene glycol hydrogel for intervertebral disc regeneration. Acta Biomater. 10, 1102–1111 (2014).

60. Seidlits, S. K. et al. Fibronectin-hyaluronic acid composite hydrogels for three-dimensional endothelial cell culture. Acta Biomater. 7, 2401–2409 (2011).

61. Lutolf, M. P. & Hubbell, J. A. Synthesis and physicochemical characterization of end-linked poly(ethylene glycol)-co-peptide hydrogels formed by Michael-type addition. Biomacromolecules 4, 713–722 (2003).

62. Cambria, E. et al. Covalent Modification of Synthetic Hydrogels with Bioactive Proteins via Sortase-Mediated Ligation. Biomacromolecules 16, 2316–2326 (2015).

63. Goldshmid, R. & Seliktar, D. Hydrogel Modulus Affects Proliferation Rate and Pluripotency of Human Mesenchymal Stem Cells Grown in Three-Dimensional Culture. ACS Biomaterials Science and Engineering 3, (2017).

64. Bott, K. et al. The effect of matrix characteristics on fibroblast proliferation in 3D gels. Biomaterials 31, 8454–8464 (2010).

65. Jones, D. R., Marchant, R. E., Von Recum, H., Gupta, A. Sen & Kottke-Marchant, K. Photoinitiator-free synthesis of endothelial cell-adhesive and enzymatically degradable hydrogels. Acta Biomater. 13, 52–60 (2015).

66. Semenza, G. L. Vasculogenesis, angiogenesis, and arteriogenesis: Mechanisms of blood vessel formation and remodeling. J. Cell. Biochem. 102, 840–847 (2007).

67. Ferrara, N. & Kerbel, R. S. Angiogenesis as a therapeutic target. Nature 438, 967–974 (2005).

68. Ribatti, D. The chick embryo chorioallantoic membrane (CAM). A multifaceted experimental model. Mech. Dev. 141, 70–77 (2016).

69. Baum, O. et al. VEGF-A Promotes Intussusceptive Angiogenesis in the Developing Chicken Chorioallantoic Membrane. 447–457 (2010). doi:10.1111/j.1549-8719.2010.00043.x

70. Marinaccio, C., Nico, B. & Ribatti, D. Differential expression of angiogenic and anti-angiogenic molecules in the chick embryo chorioallantoic membrane and selected organs during embryonic development. 916, 907–916 (2014).

71. Shekaran, A. et al. Bone regeneration using an alpha 2 beta 1 integrin-specific hydrogel as a BMP-2 delivery vehicle. Biomaterials 35, 5453–5461 (2014).

72. Fernandez de Grado, G. et al. Bone substitutes: a review of their characteristics, clinical use, and perspectives for large bone defects management. Journal of Tissue Engineering (2018). doi:10.1177/2041731418776819

