## Supplementary Material for "Engineered full-length Fibronectin-based hydrogels sequester and present growth factors to promote regenerative responses in vitro and in vivo" (from another version of this preprint: version 1, 2019-06-29)

### Supplementary Methods

#### Enzyme-linked immunosorbent assay

Enzyme-linked immunosorbent assay (ELISA) plates were coated with the capture antibody rabbit-polyclonal-anti-FN (Sigma) overnight at RT. Then, the capture antibody was washed three times with washing buffer (0.05% Tween 20 in PBS). After washing, the plate was blocked using blocking buffer (1% BSA) for 1 h at RT. After blocking, the plate was washed three times using washing buffer. Then, samples (PEGylated fibronectin or native fibronectin) were added at different concentrations to the plate for 1 h at RT. After sample incubation, the plate was washed three times with washing buffer. Then, the monoclonal anti-FN primary antibodies (either HFN7.1 (1:330, Developmental Studies Hybridoma Bank), P5F3 (1:2000, Santa Cruz Biotechnology) or MAB1892 (1:1000, Millipore)) were added for 1 h at RT. After incubation, samples were washed three times using washing buffer. Then, secondary antibody (goat-anti-mouse-horse radish peroxidase (HRP) (1:10000, Invitrogen)) was added for 1 h at RT and was washed three times with washing buffer. After that, substrate solution (1:1 V/V Reagent A:Reagent B, R&D Systems) was added for 20 min. The reaction was stopped using 4 N sulphuric acid (Stop solution, R&D Systems). The ELISA plate was read at 450 nm and 540 nm using a plate reader (BIOTEK).

#### Atomic force microscopy (AFM) imaging

Proteins (PEGylated fibronectin or native fibronectin) were adsorbed onto poly(ethyl) Acrylate (PEA) spin coated glass coverslips for 10 min at 20 µg/mL and RT as previously described[1]. AFM in AC mode (Nanowizard-3, JPK) was used to obtain phase and height images of the fibronectin-coated PEA samples. A pyramidal tip (silicon nitride, MPP-21220, $f_0$ =59-69 Hz, Bruker) was used. Images were processed using the JPKSPM data processing software.

**Cell culture**

C2C12 murine cells (Sigma, passage 4-7) were used for early adhesion studies and viability in PEG hydrogels. C2C12s were grown in DMEM high glucose without pyruvate and 20% FBS.

**Cell adhesion on PEGylated fibronectin**

PEGylated and native fibronectin were adsorbed onto glass coverslips at 20 µg/mL for 1 h at RT. After adsorption, samples were washed twice with PBS and kept in the incubator (37°C, 5% $CO_2$) until seeding. C2C12 cells were seeded onto the glass coverslips at 10,000 cells/$cm^2$ using FBS free media. Cells were let to attach for 3 h (37°C and 5% CO2). After that time, cells were fixed with 4% paraformaldehyde for 30 min at RT and kept at 4°C.

**Polyacrylamide gel electrophoresis**

The polyacrylamide gel electrophoresis (PAGE) was run using 0.1% SDS in a Trys-Glycine running buffer (Bio-Rad). Samples were loaded using a reducing loading buffer prepared by mixing Laemmli buffer (4X, Bio-Rad) and β-mercaptoethanol (350 mM, Bio-Rad), following manufacturer's instructions. Fifteen microliters of each sample (sample + loading buffer) were loaded and 5 µL of the marker (colour pre-stained protein ladder, broad range (11-245 kDa, New England Biolabs) were used. A gradient precast gel (BioRad, 4-20% TGX-PAGE) was used to run the samples. The running conditions were 100 V, 1800 min. Gels were incubated in fixing solution (methanol, water and glacial acetic acid 45:45:10 (V/V/V)) for 30 min at RT. Then, gels were stained with 0.025% Coomassie Blue G250 (Bio-Rad) in 10% acetic acid overnight at RT using an orbital shaker. Finally, each gel was destained twice in 10% acetic acid and transferred to water. Images of the gel were taken using the scanner of a canon image runner advance C5235i multifunctional laser printer. Each condition was run in triplicate in the same gel except for the protein ladder, which was run twice in the same gel.

**Image analysis**

**Cell morphology analysis**

Cell shape descriptors were measured using ImageJ 1.51v (National Institutes of Health, US). Briefly, actin cytoskeleton images were binarized using a threshold function. Then, the wand tracing tool was used to select the outline of the cell and the measure function was used to calculate parameters such as cell area, aspect ratio and roundness. The aspect ratio is defined as the ratio between the major and minor axis of the shape selected and the roundness was calculated as follows: $Roundness=((4*area))/((\pi*(major\ axis)^2))$

**Focal adhesion (FA) analysis**

Vinculin images were used for the FA analysis. They were uploaded to the Focal Adhesion Analysis Server[2] along with the corresponding actin cytoskeleton image to use as a cell mask. The results obtained include a binarized image of the FAs and measurements for several parameters, from which the FA number per cell, the FA area and size were used.

### Supplementary References

1. Llopis-hernández, V. *et al.* Material-driven fibronectin assembly for high-efficiency presentation of growth factors. 1–10 (2016).

2. Berginski, M. E. & Gomez, S. M. The Focal Adhesion Analysis Server: a web tool for analyzing focal adhesion dynamics. *F1000Research* (2013). doi:10.12688/f1000research.2-68.v1

### Supplementary Figures

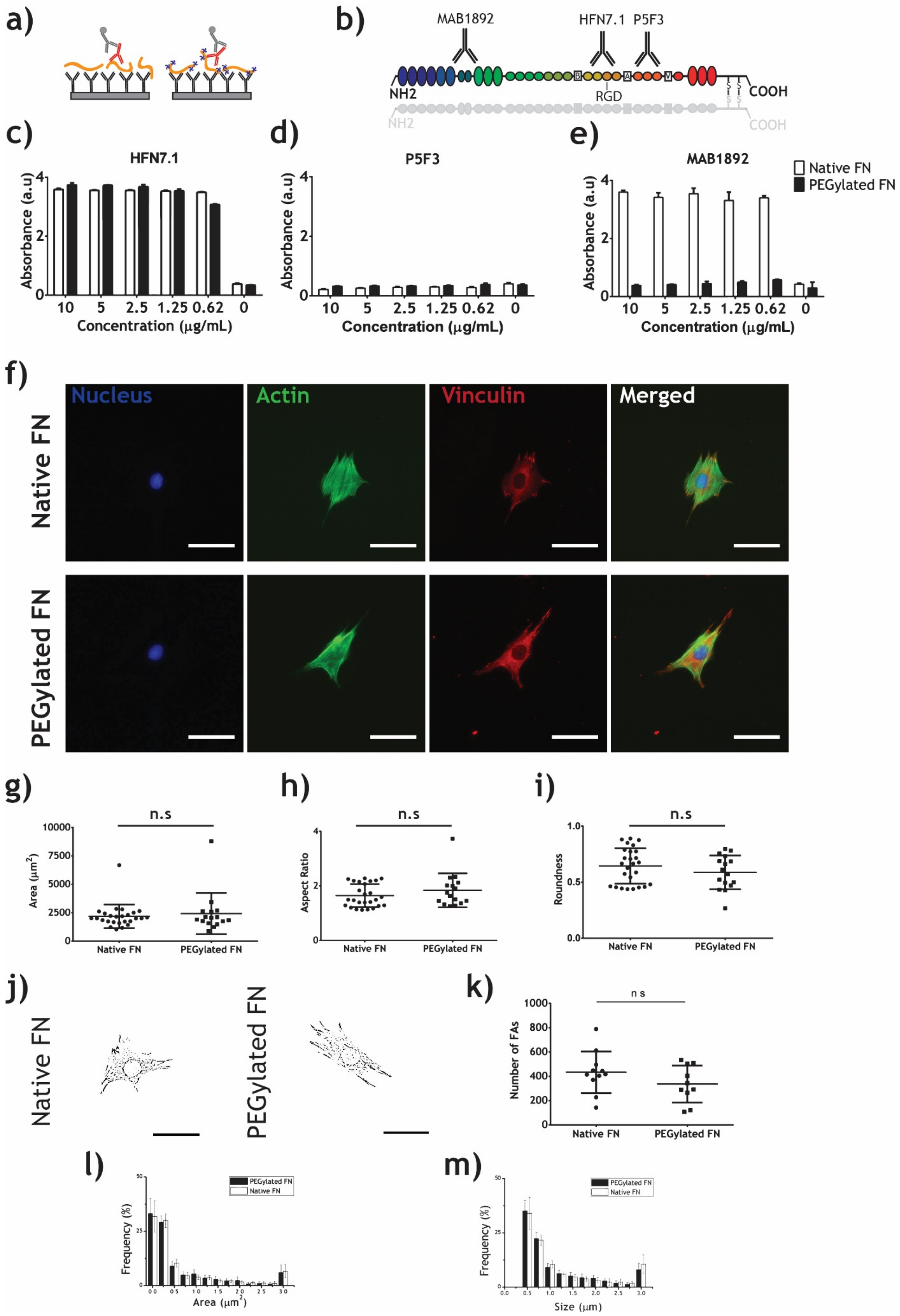

**Figure S1. PEGylated fibronectin maintains biological activity of key domains.**

**(a)** Sketch of sandwich ELISA performed for native and PEGylated fibronectin in solution, (**b**) scheme of fibronectin molecule and domains targeted using monoclonal antibodies HFN7.1, P5F3 and MAB 1892 for ELISA assays. Absorbance values for different concentrations of native and PEGylated fibronectin when using **(c)** HFN7.1, **(d)** P5F3 and **(e)** MAB 1892 antibodies. **(f)** Representative fluorescence images (nucleus in blue, actin in green and vinculin in red) of C2C12 cells seeded on top of native or PEGylated fibronectin for 3h (scale bar: 50 µm). Graphs showing **(g)** cell area, **(h)** cell aspect ratio and **(i)** cell roundness for cells seeded on top of native or PEGylated fibronectin. **(j)** Representative images of segmented focal adhesions (FAs) of C2C12s seeded on native or PEGylated fibronectin for 3h (scale bar: 50 µm). Graphs showing **(k)** number of focal adhesions, **(l)** frequency of FA area and **(m)** size.

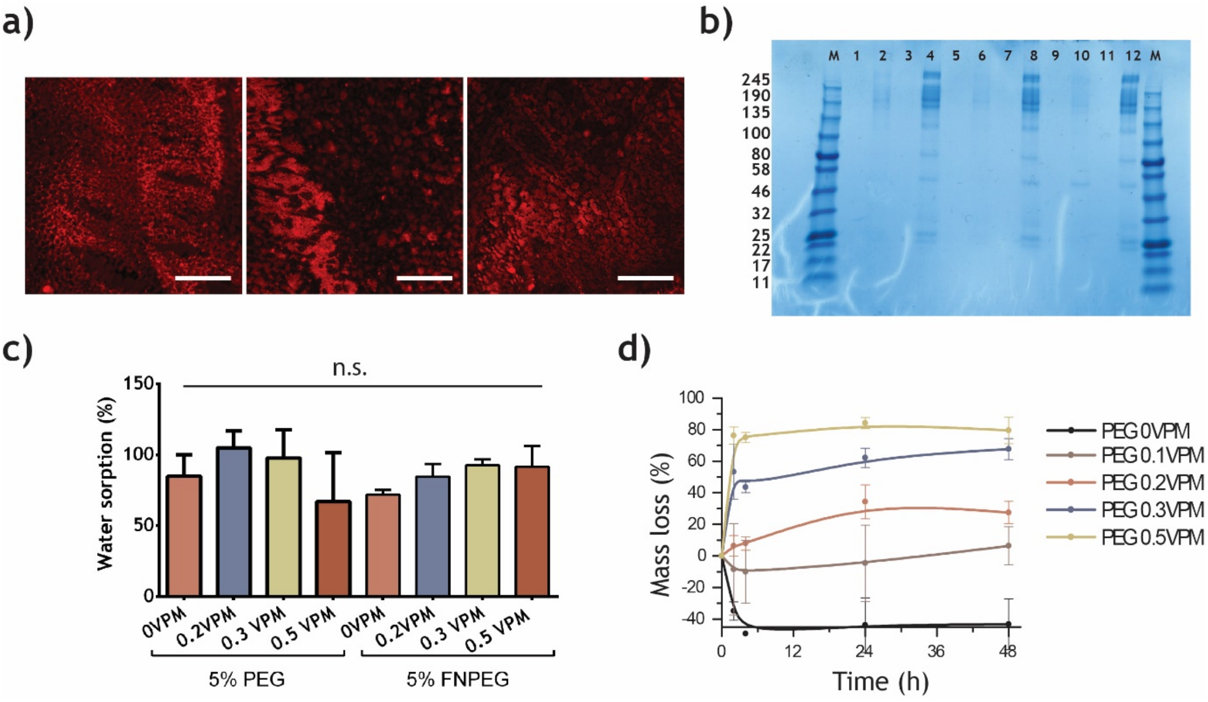


**Figure S2. Fibronectin hydrogels characterisation.**

**(a)** immunofluorescence of fibronectin (in red) on cryosections of 3, 5 and 10 wt. % hydrogels (from left to right, scale bar: 100 µm); **(b)** supernatants from fibronectin release were run in a SDS-PAGE and were stained with Coomassie blue. **(c)** Water absorbance of 5 wt.% PEG only and FNPEG hydrogels with different ratios of degradable crosslinker (0, 0.2, 0.3, 0.5 VPM peptide); **(d)** Degradation profile of PEG only hydrogels with different ratios of degradable crosslinker.

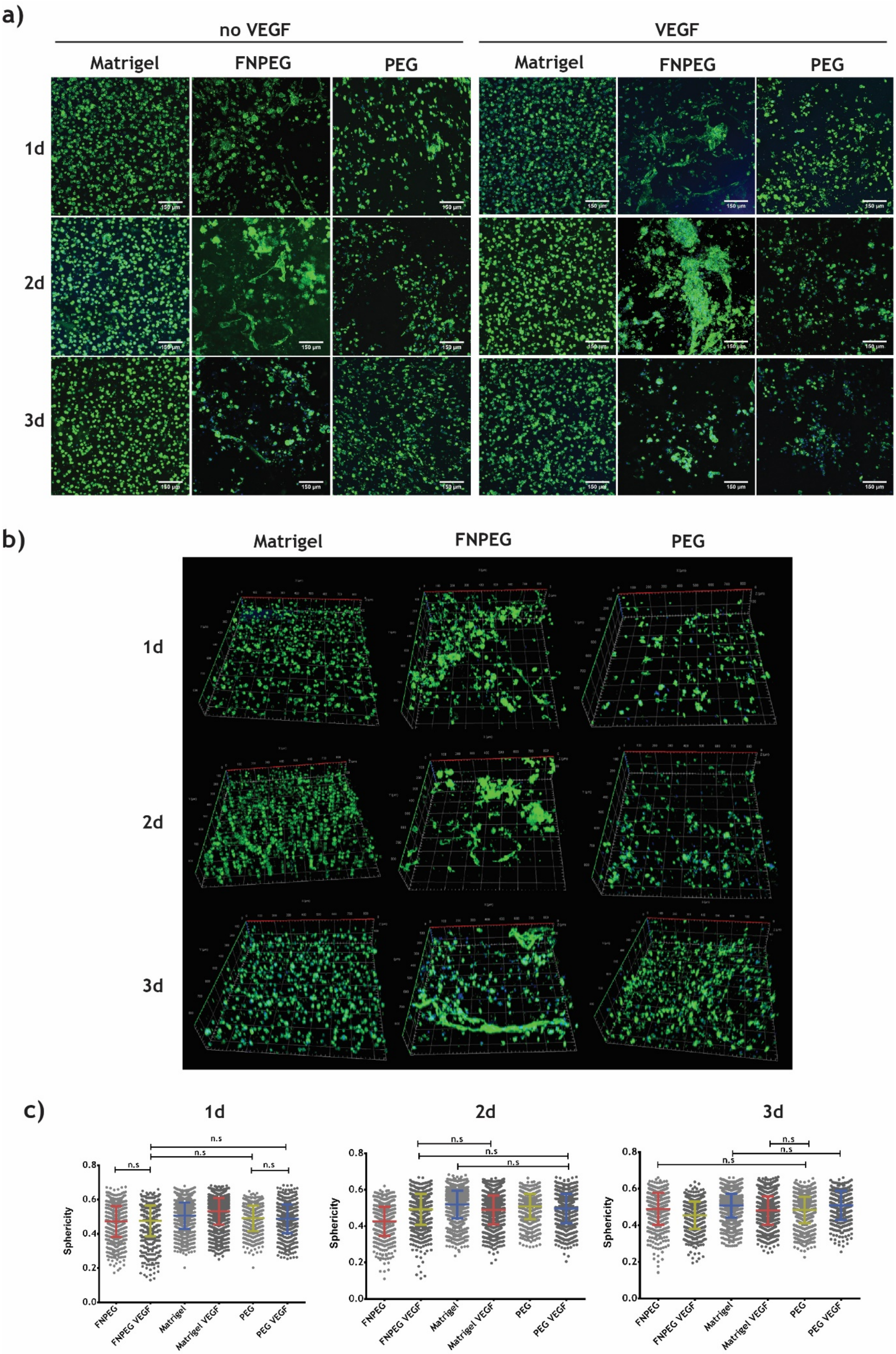
a)
no VEGF
VEGF
Matrigel
FNPEG
PEG
Matrigel
FNPEG
PEG
1d
2d
3d
b)
Matrigel
FNPEG
PEG
1d
2d
3d
c)
1d
2d
3d
Sphericity
n.s
FNPEG
FNPEG VEGF
Matrigel
Matrigel VEGF
PEG
PEG VEGF

**Figure S3. Fibronectin hydrogels promote microvasculature growth in 3D.**

**(a)** Panel of maximum Z-axis projections of HUVECs within different conditions tested at different timepoints (nucleus in blue and actin in green, scale bar: 150 µm). **(b)** Representative 3D reconstructions of HUVECs within Matrigel, FNPEG or PEG at days 1, 2 and 3 (without VEGF, nucleus in blue, actin in green). **(c)** Graphs showing sphericity values at days 1, 2 and 3 for different conditions tested.

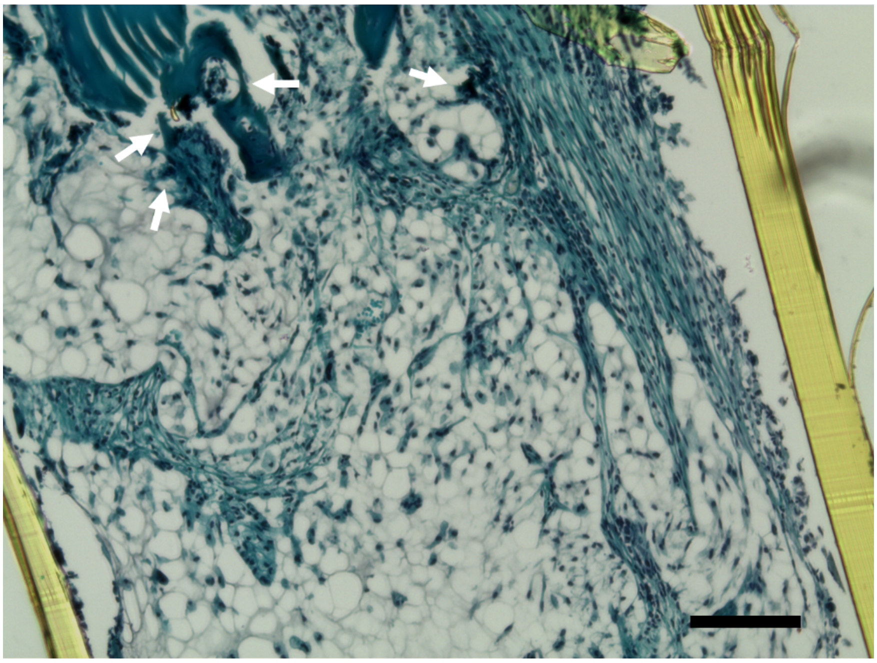

**Figure S4. Fibronectin hydrogels promote bone growth in vivo.**

Histological section of FNPEG-0 condition stained with safranin O, fast green and haematoxylin. Resorption bays are marked by arrows. Scale bar: 100 um.
